# Towards *Mycobacterium tuberculosis* detection at the point-of-care: a brighter solvatochromic probe permits the detection of mycobacteria within minutes

**DOI:** 10.1101/2020.05.29.124008

**Authors:** Mireille Kamariza, Samantha G. L. Keyser, Ashley Utz, Benjamin D. Knapp, Green Ahn, C. J. Cambier, Teresia Chen, Kerwyn Casey Huang, Carolyn R. Bertozzi

## Abstract

There is an urgent need for point-of-care tuberculosis (TB) diagnostic methods that are fast, inexpensive, and operationally simple. Here, we report on a bright solvatochromic dye trehalose conjugate that specifically detects *Mycobacterium tuberculosis* (Mtb) in minutes. 3-hydroxychromone (3HC) dyes, known to yield high fluorescence quantum yields, exhibit shifts in fluorescence intensity in response to changes in environmental polarity. We synthesized two analogs of 3HC-trehalose conjugates (3HC-2-Tre and 3HC-3-Tre) and determined that 3HC-3-Tre has exceptionally favorable properties for Mtb detection. 3HC-3-Tre-labeled mycobacterial cells displayed a 10-fold increase in fluorescence intensity compared to our previously reports on the dye 4,4-*N,N*-dimethylaminonapthalimide (DMN-Tre). Excitingly, we detected fluorescent Mtb cells within 10 minutes of probe treatment. Thus, 3HC-3-Tre permits rapid visualization of mycobacteria that ultimately could empower improved Mtb detection at the point-of-care in low-resource settings.

## INTRODUCTION

With 1.2 million deaths and 10 million new cases in 2018, tuberculosis (TB) is the most lethal infectious disease in the world (1). Early detection of the bacterium *Mycobacterium tuberculosis* (Mtb), the causative agent of TB, followed by appropriate treatment could prevent most deaths (2). The gold standard for TB diagnosis remains a labor-intensive culture test that requires weeks of incubation time in specialized facilities. Although more rapid tests are available, they present several important limitations. PCR-based tests are expensive and require skilled technicians. Microscopy-based methods are attractive in low-resource settings as they are low-cost, have fast turnaround times, and report on people at greatest risk of transmission and death (3). As a result, the sputum smear microscopy test is the most widely used technique for TB diagnosis. The century-old smear test is based on the susceptibility of fluorescent auramine dye or colored Ziehl-Neelson (ZN) stain to accumulate within the highly hydrophobic mycobacterial cell wall (4–7). While effective for identification of Mtb cells, this process requires multiple wash steps to reduce non-specific background fluorescence, or in the case of the ZN test, a rigorous counterstaining procedure so that stained Mtb cells can be visualized (8,9). Moreover, the smear test does not distinguish live from dead cells, and this capability is vital in order to assess treatment efficacy early and accurately (2).

In the last decade, we and others have leveraged the trehalose metabolism of mycobacteria to mark them for detection by various imaging methods (10–20). Exogenous trehalose molecules can be directly mycolylated at the 6 position by antigen 85 (Ag85) enzymes to form trehalose monomycolates (TMM) that are inserted into the mycobacterial cell wall, termed the mycomembrane (10). Researchers have shown that Ag85 enzymes are promiscuous enough to tolerate perturbations of varying sizes such as azide (11), alkyne (12), fluorine (13–15), and fluorophore (13,16) groups that permit visualization of the mycomembrane as long as the cell is metabolically active. However, these fluorescent probes require extensive washing before imaging in order to reduce background signal.

Fluorogenic probes, i.e. probes that turn on when metabolized in cells, have proven better suited for TB detection as they require minimal processing. Previous studies have used quenched trehalose fluorophores that become unquenched by Ag85 activity, allowing visualization of growing mycobacterial cells in real-time (17). A dual enzyme-targeting fluorogenic probe allowed the detection of Mtb cells within an hour using a microfluidic system (18). In addition, we reported on a solvatochromic trehalose probe (DMN-Tre) that can detect live Mtb cells in TB patient sputum samples (19). Solvatochromic dyes change their color or fluorescence intensity based on the polarity of the solvent. As a result, these compounds are advantageous for monitoring changes in hydrophobicity of a molecule of interest (20). Upon acylation of DMN-Tre by Ag85 and insertion of the corresponding trehalose monomycolate (TMM) analog into the mycomembrane, dye fluorescence is turned on, which allows the detection of live mycobacteria with a fluorescence microscope (**Figure 1A**) (19,21). In this context, DMN-Tre has many favorable properties for point-of-care deployment: its operationally simple procedure does not require any wash steps, and it is synthetically convenient and chemically stable. However, the DMN dye is a fluorophore of relatively low brightness, with a molar extinction coefficient of 8800 M^-1^ cm^-1^ in TBS buffer (20) and a low quantum yield of fluorescence in low-dielectric solvents (0.288 in DMF) (22). Consequently, in practice we found that: 1) low-powered fluorescence microscopes currently available in TB health centers may miss labeled cells, and 2) Mtb cells must be labeled with DMN-Tre for at least one hour to achieve detectable levels of fluorescence using standard clinical fluorescence microscopes.

**Figure 1.**
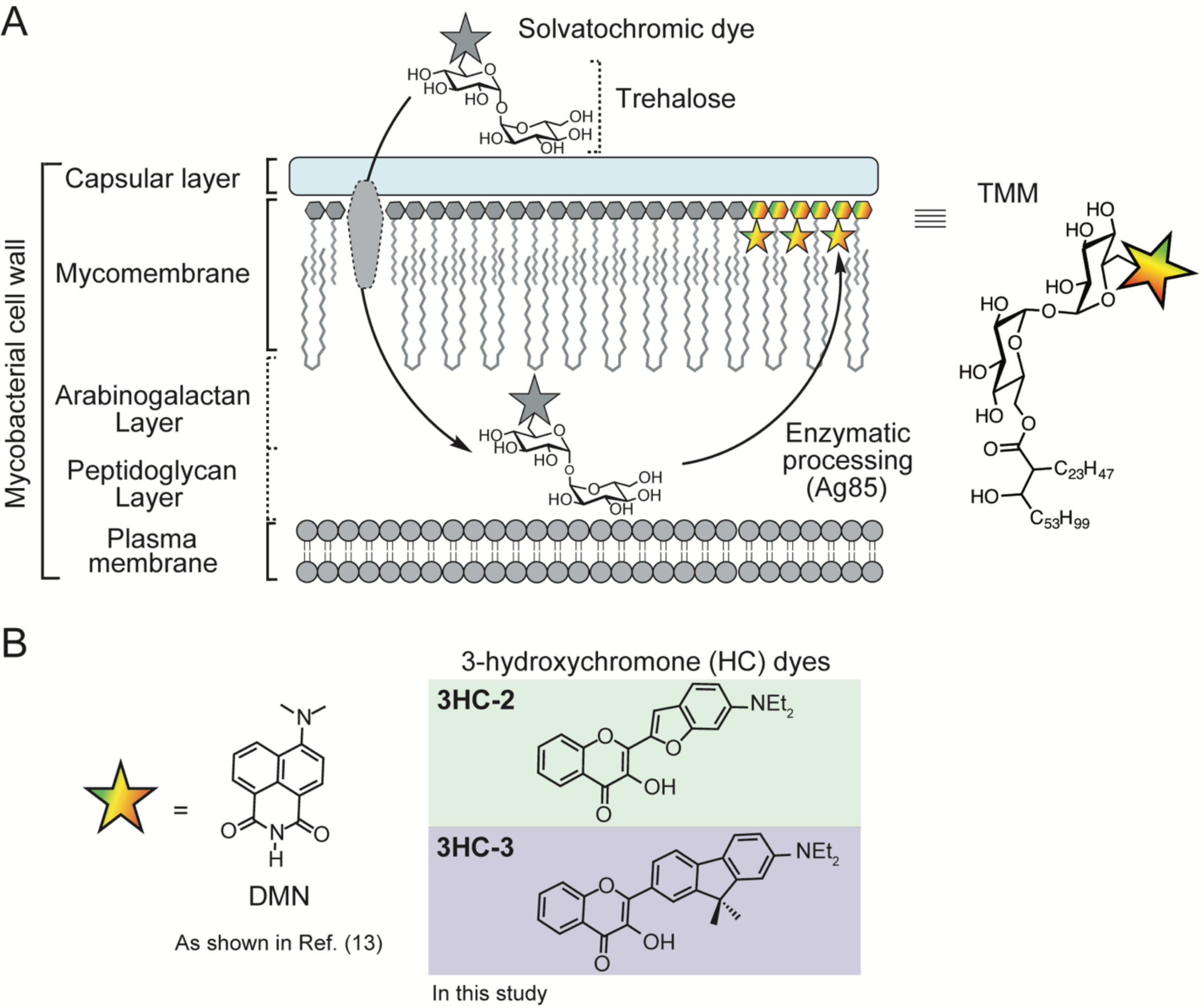
Solvatochromic trehalose probes label the mycobacterial mycomembrane. **(A)** Solvatochromic trehalose probes are converted by mycobacteria to the corresponding trehalose monomycolate (TMM, structure on right) analogs and inserted into the mycomembrane. There, they undergo fluorescence turn-on, enabling detection of labeled cells by fluorescence microscopy. **(B)** Chemical structures of solvatochromic dyes described in this study.

To address these obstacles for point-of-care TB detection, we aimed to develop probes that are brighter and enable faster detection of live Mtb cells. Here, we report the development of a brighter solvatochromic trehalose probe, based on the 3-hydroxychromone (3HC) dye (**Figure 1B**). The 3HC trehalose conjugate (termed 3HC-3-Tre) demonstrated high fluorescence turn-on in hydrophobic solvents as well as when incorporated in the mycobacterial cell surface. This labeling was specific to the trehalose moiety and was detectable without any wash steps. Additionally, the fluorescence intensity of 3HC-3-Tre was 10-fold brighter than DMN-Tre. Finally, the high signal-to-noise ratio of 3HC-3-Tre permitted simple detection of labeled Mtb cells within 10 minutes. Thus, 3HC-3-Tre reagent permits the rapid visualization of mycobacteria and ultimately, could be used to improve Mtb detection at the point-of-care in low-resources settings.

## RESULTS

### Synthesis of 3HC solvatochromic dyes bound to trehalose

To design a probe with stronger turn-on fluorescence than DMN, we relied on previously reported solvatochromic dyes that fit our target profile. We identified a highly promising class of solvatochromic dyes that are well characterized, synthetically tractable, and have been used in living systems with minimal perturbations (**Figure 1B**) (20). These dyes are based on a 3-hydroxychromone (3HC) scaffold with the advantage of greater tunability due to their synthetic modularity and high quantum fluorescence yield. Moreover, they are further red-shifted, which may minimize background fluorescence.

We began by synthesizing 6-Br-Ac_7_-Tre (**Figure 2**). An Appel reaction using N-bromosuccinimide and triphenylphosphine resulted in a mixture of mono-brominated target compound (6-Br-Tre), a dibrominated side product (6,6’-dibromo-6,6’-dideoxy-α,α’-trehalose), and unreacted starting material. The crude material was then acetylated and purified to give 6-Br-Ac_7_-Tre (**Compound 2**). Next, we considered the synthesis of 3HC dyes, here referred to as 3HC-3 (**Compound 3**) and 3HC-2 (**Compound 4**) (23–26). We followed a previous study (23) to achieve synthesis of the aldehyde precursor to 3HC-2 by reacting 3-diethylaminophenol with bromoacetaldehyde diethyl acetal (**Scheme S1**). The intermediate was purified, then subsequently treated with phosphorus(V) oxychloride and *N,N*-dimethylformamide (DMF) to form the benzofuran and to install an aldehyde at the 2-position via a Vilsmeier-Haack reaction. To synthesize the aldehyde precursor to 3HC-3 (**Compound 3**), we nitrated 2-bromo-9,9-dimethylfluorene at the 7-position, reduced the nitrate to an amine, alkylated the amine using ethyl iodide, and converted the bromine to an aldehyde through a Bouveault reaction. To form 3HC-2 and 3HC-3, each aldehyde was reacted with 2’-hydroxyacetophenone, then treated with hydrogen peroxide to obtain the desired products (**Figure 2B**) (27). Finally, to create the dye-Tre probes, the aromatic hydroxyl group on the dye was deprotonated with potassium carbonate and used to displace the bromine atom on 6-Br-Ac_7_-Tre (**Figure 2C**) (28). The crude dye-Ac_7_-Tre was then deacetylated with catalytic sodium methoxide and purified to produce the final dye-Tre products in yields ranging from 9-63% over three steps. We also synthesized glucose control compounds in a similar fashion (**Scheme S2**).

**Figure 2.**
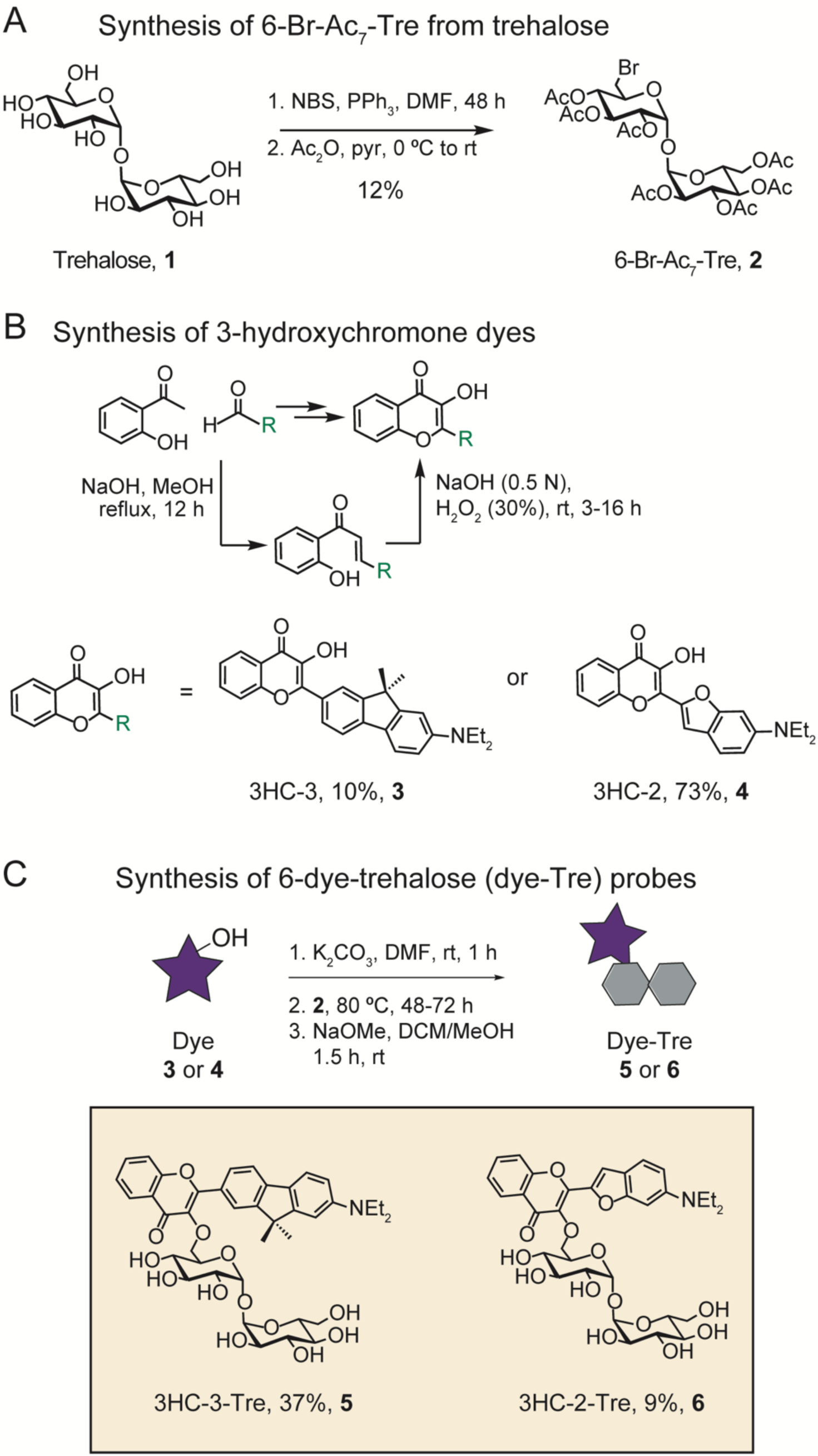
Synthetic scheme for 3-hydroxychromone (3HC) trehalose (Tre) dye conjugates.

### *3HC-trehalose probes label* Mycobacterium smegmatis *(Msmeg) cells more effectively than DMN-Tre*

With these probes in hand, we proceeded to characterize their fluorescence intensities and emission spectra in mixtures with various ratios of dioxane and water. The dyes were excited at the optimal excitation wavelengths (405 nm for DMN-Tre and 3HC-3-Tre, 488 nm for 3HC-2-Tre) and fluorescence intensities were measured over a range of emission wavelengths (**Figure 3**). As expected, all probes displayed increased fluorescence as the amount of dioxane increased. DMN-Tre and 3HC-3-Tre both absorbed most strongly at 405 nm and had comparable fluorescence intensities in each solvent mixture tested, although the spectra for 3HC-3-Tre were slightly red-shifted (**Figure 3A,B**). 3HC-2-Tre responded well to excitation at 405 nm and 488 nm, although its fluorescence intensity was less sensitive to changes in hydrophobicity overall (**Figure 3C**). However, because the emission spectra underwent a bathochromic shift as solvent polarity increased, significant differences in intensity still occurred between 500 and 550 nm, the approximate range of wavelengths allowed through the GFP emission filter.

**Figure 3.**
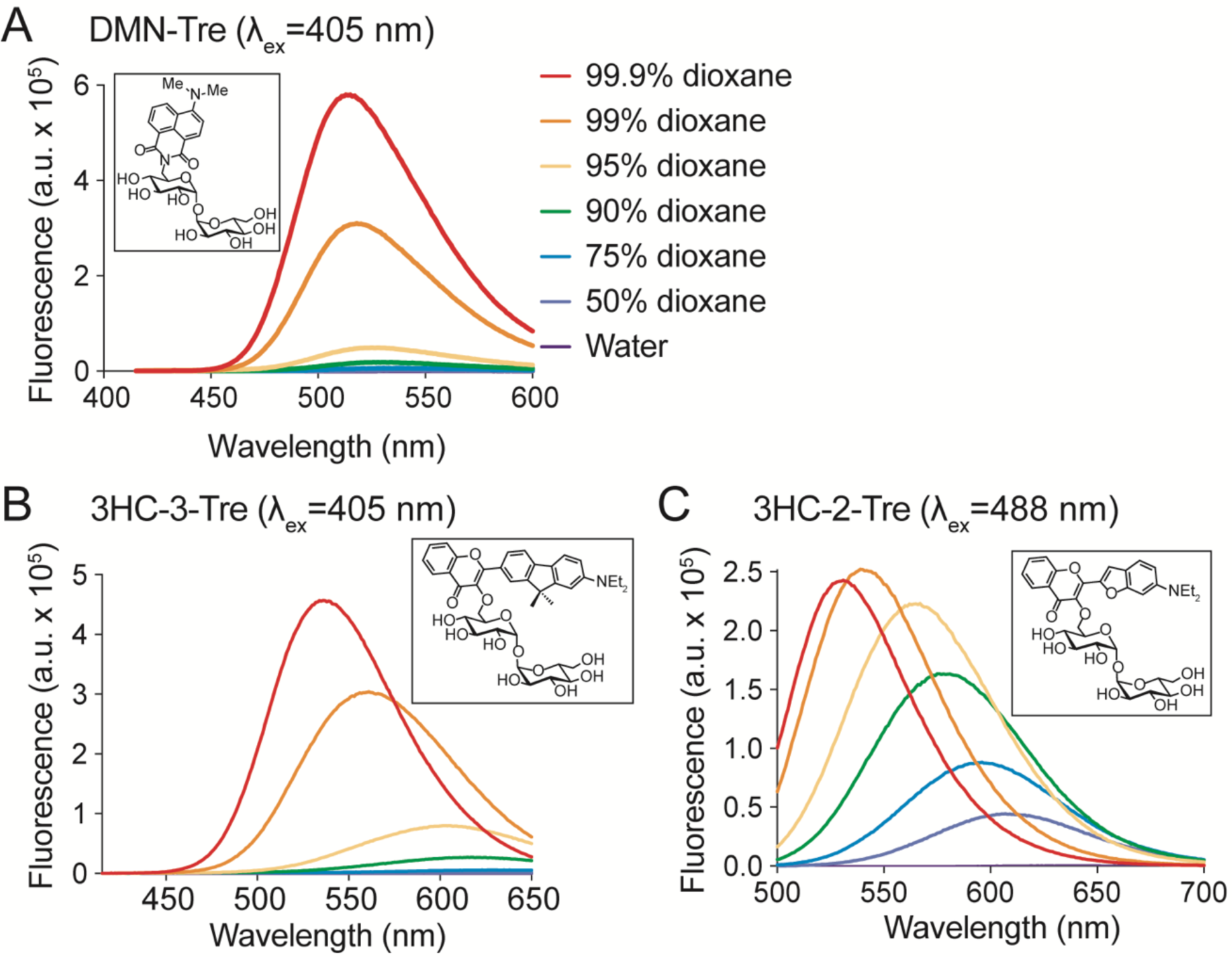
Emission spectra of 3-hydroxychromone trehalose dyes. Fluorescence spectra of (**A**) DMN-Tre (ex. 405 nm), (**B**) 3HC-3-Tre (ex. 405 nm) and (**C**) 3-HC-2-Tre (ex. 488 nm) in solvent systems with the indicated ratios of dioxane in water.

We also assessed labeling conditions for *Mycobacterium smegmatis* (Msmeg), a nonpathogenic and fast-growing member of the *Mycobacterium* genus commonly used as a model organism for Mtb. Msmeg cells were grown to an optical density at wavelength 600 nm (OD_600_) of 0.5, then incubated with 1, 10, or 100 μM of each probe for 1 hour at 37 °C, washed three times, and analyzed by flow cytometry using a variety of excitation and emission filter sets (**Figure 4**). As expected, we observed that all trehalose probes labeled Msmeg in a concentration-dependent manner. As well, 3HC-3-Tre and 3HC-2-Tre-labeled cells reached much higher levels of fluorescence intensity compared to DMN-Tre-labeled bacteria (approximately 10-fold and 100-fold higher for 3HC-3-Tre and 3HC-2-Tre, respectively). While DMN-Tre’s fluorescence was optimally detected with 405/525 (ex/em) nm filter sets (**Figure 4A**), we determined that the optimal fluorescence detection filter sets for 3HC-3-Tre and 3HC-2-Tre probes were 405/525 and 488/525 nm, respectively (**Figure 4B,C**). Excitingly, even with 10-fold lower concentrations, 3HC-3-Tre-labeled Msmeg cells demonstrated nearly 5-fold greater fluorescence intensity compared to DMN-Tre.

**Figure 4.**
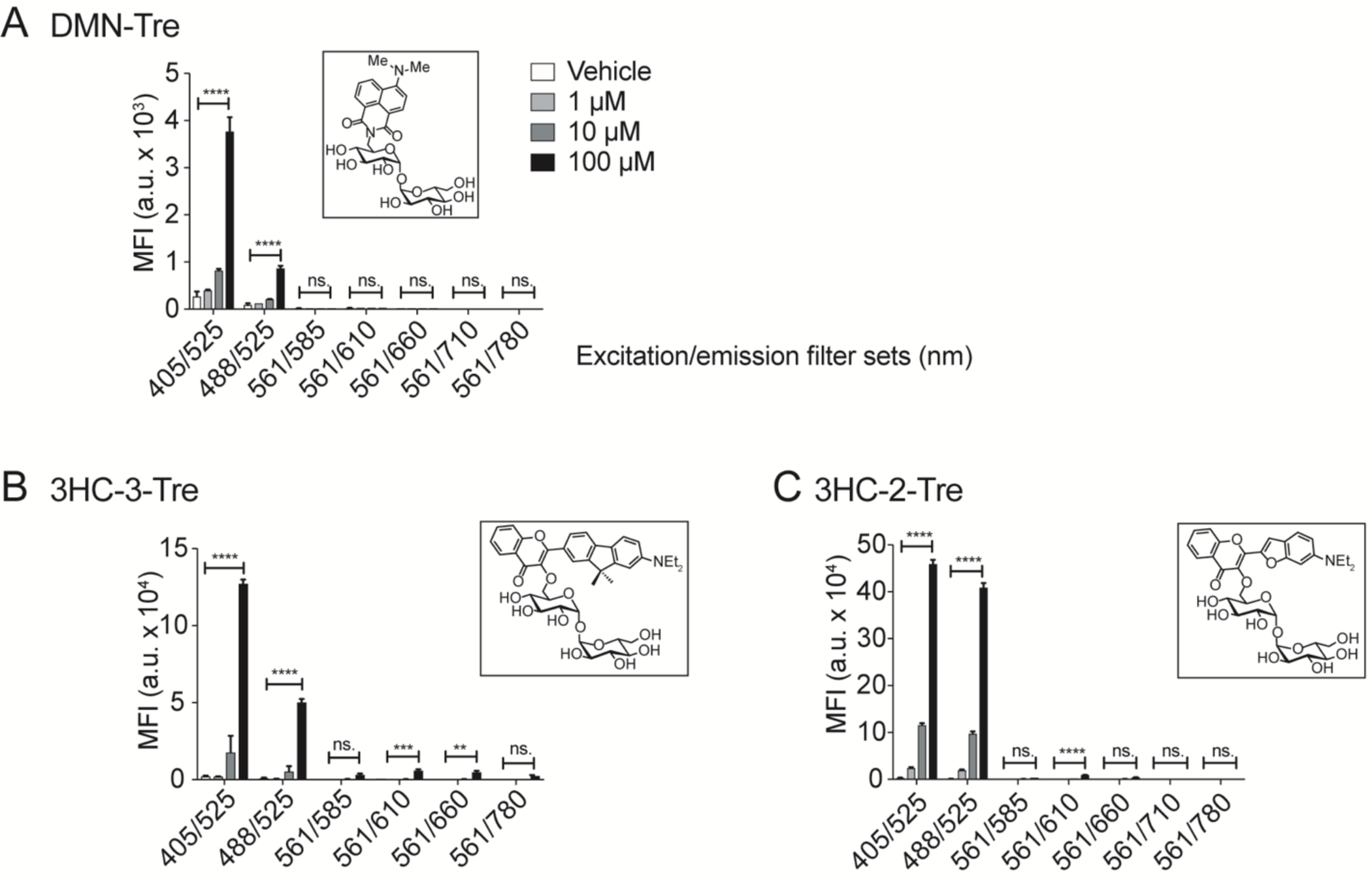
Flow cytometry analysis of Msmeg cells labeled with solvatochromic trehalose dyes using various excitation and emission filter sets. Flow cytometry analysis of Msmeg labeled with (**A**) DMN-Tre, (**B**) 3HC-3-Tre or (**C**) 3HC-2-Tre. All dyes showed increased labeling at the highest concentration (100 µM). Cells at OD_600_=0.5 were incubated with the indicated dye-trehalose probe concentrations for 1 h at 37 °C. MFI = Mean fluorescence intensity. Data are means ± SEM from at least two independent experiments. Data were analyzed by two-way analysis of variance (ANOVA) test (*: *p*<0.05, **: *p*<0.01, ***: *p*<0.001, ****: *p*<0.0001, ns: not significant).

### 3HC-3-Tre rapidly and stably labels mycobacteria in a trehalose-dependent manner

For the dyes to be useful in the field, the labeling procedure must follow a simple protocol. Thus, we sought to determine whether a wash step is necessary based on the degree of background signal when Msmeg cells are labeled with each trehalose probe, glucose-dye control, or free dye. We incubated Msmeg cells with DMN-Tre, 3HC-3-Tre, or 3HC-2-Tre (along with their respective glucose and sugar-free analogs) for 1 hour at 37 °C and performed fluorescence microscopy either immediately or after a wash step with PBS (**Figure 5**). As anticipated, we observed fluorescent Msmeg cells labeled with DMN-Tre, while the fluorescence of DMN-Glc-labeled Msmeg was washed off with PBS (**Figure 5A**), suggesting that the non-specific turn-on of DMN-Glc molecules is likely due to proximity to Msmeg cells. Surprisingly, we observed no fluorescence with 3HC-3-Glc-labeled Msmeg cells, even in unwashed samples (**Figure 5B, Figure S1A**), suggesting that internalization of the probe is required for fluorescence turn-on of 3HC-3. Interestingly, we observed labeling of Msmeg cells with 3HC-2 and 3HC-2-Glc, even after washing (**Figure 5C, Figure S1B**), suggesting that 3HC-2 labeling is not specific to the trehalose pathway. Intrigued by this result, we wondered whether the fluorescence labeling of Msmeg from 3HC-2 dye conjugates would reflect the known trehalose insertion patterning.

**Figure 5.**
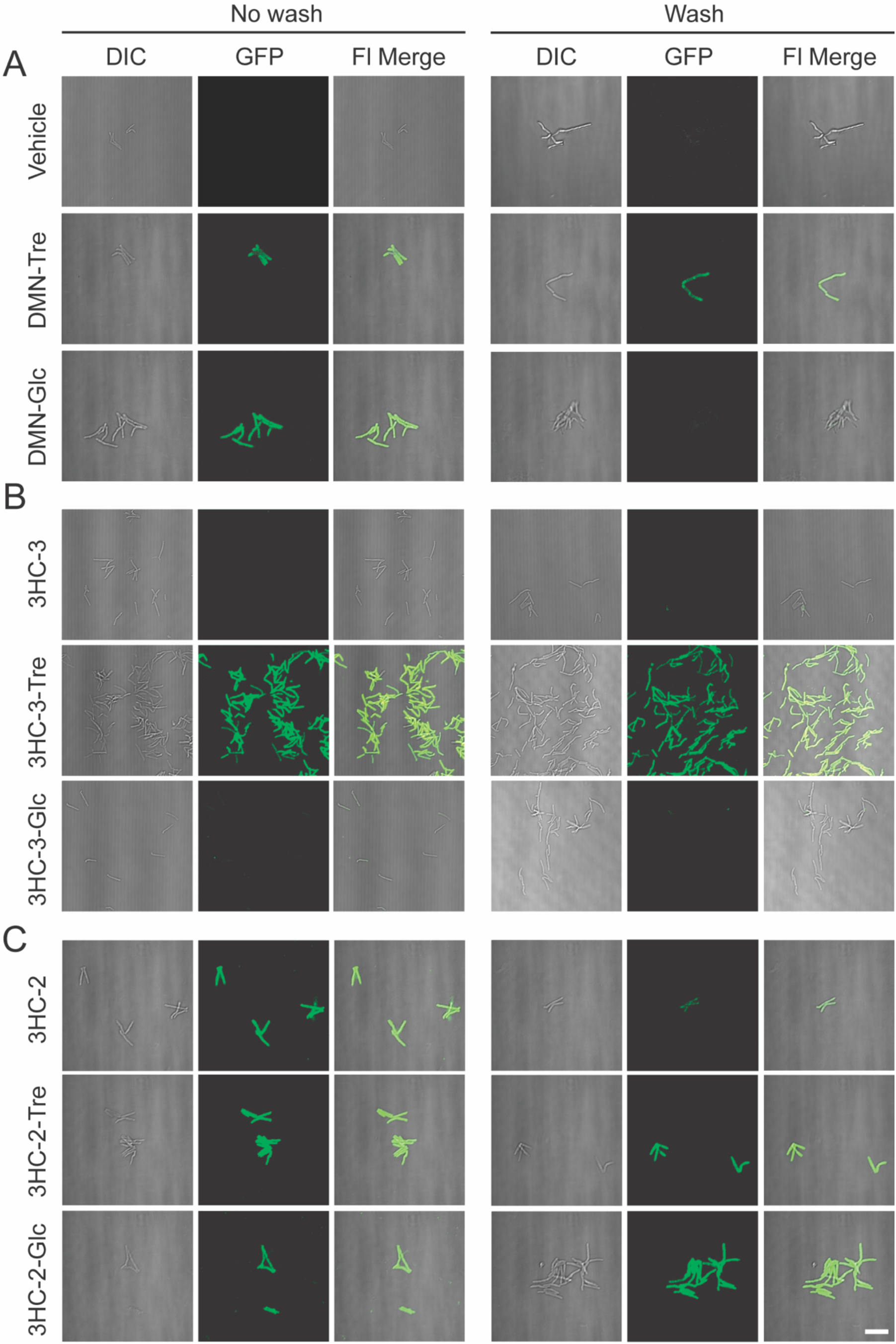
Unlike 3HC-2-Tre, 3HC-3-Tre labeling is dependent on the trehalose moeity. Epifluorescence microscopy of Msmeg cells treated with (**A**) 100 µM DMN-Tre or DMN-Glc or no dye control (Vehicle); (**B**) 100 µM of 3HC-3, 3HC-3-Tre, or 3HC-3-Glc; (**C**) 100 µM of 3HC-2, 3HC-2-Tre, or 3HC-2-Glc. 3HC-3-Tre showed the most efficient labeling of Msmeg. Cells were incubated with the indicated dyes for 1 h at 37 °C. Cells were smeared directly (No wash) or washed 3 times with PBS then smeared onto a microscope slide (Wash). Scale bar: 10 µm.

Previous studies demonstrated that exogenous trehalose molecules are mycolylated at the 6 position via action of Ag85 enzymes, which are localized at the septa and poles of the cell envelope (10,19). We hypothesized that 3HC-2 labeling occurs in an Ag85-independent manner and therefore would not exhibit polar and septal fluorescence. Using total internal reflection fluorescence (TIRF) microscopy, we placed Msmeg cells into a microfluidic flow cell (Methods), introduced liquid growth medium containing DMN-Tre, 3HC-3-Tre, 3HC-2-Tre, or 3HC-2-Glc, and performed time-lapse microscopy (**Figure 6, Figure S2**). Similar to DMN-Tre, 3HC-3-Tre labeling of Msmeg cells showed septal and polar labeling (**Figure 6A**, top two panels). We observed no enhancement in polar or septal fluorescence localization for Msmeg cells labeled with 3HC-2-Tre or 3HC-2-Glc (**Figure 6A**, bottom two panels), further suggesting that this labeling likely does not depend on the trehalose metabolic pathway.

**Figure 6.**
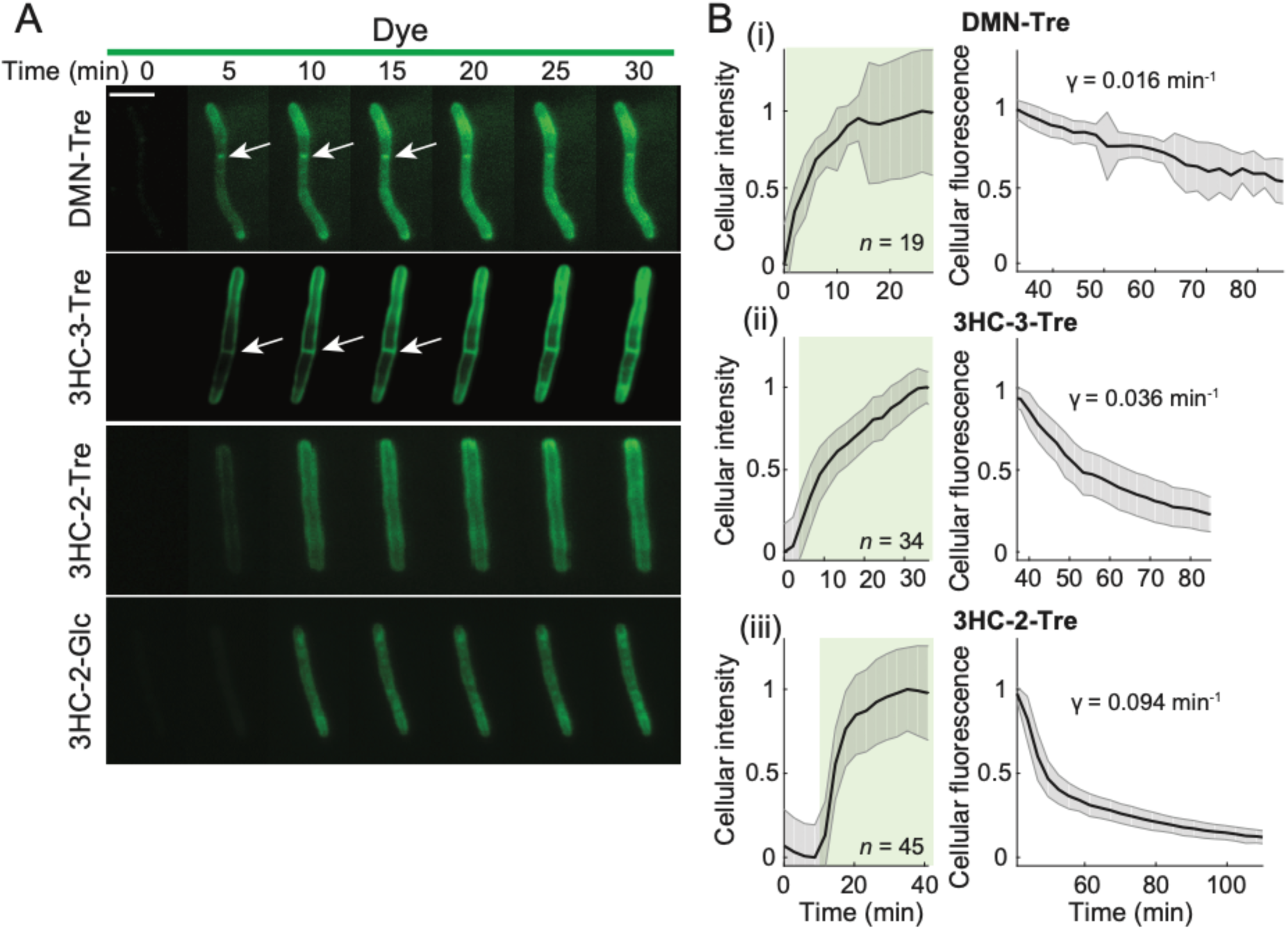
Unlike 3HC-2 dye conjugates, 3HC-3-Tre labeling is initially localized at the septum and poles. (**A**) Time-lapse microscopy of Msmeg cells treated with 100 µM DMN-Tre, 3HC-3-Tre, 3HC-2-Tre, or 3HC-2-Glc for 30 min revealed concentration of 3HC-3-Tre at cell septa and poles. White arrows denote septal labeling. Scale bar: 5 µm. (**B**) Quantitation of Msmeg fluorescence in the presence of 100 µM (i) DMN-Tre, (ii) 3HC-3-Tre, or (iii) 3HC-2-Tre during labeling for 30 min (left, volume-normalized intensity) and subsequent washing with growth medium for 1 h (right, total fluorescence). The number of cells included in each analysis (*n*) is provided in each panel. Shaded error bars represent ± 1 standard deviation.

After 30 minutes of labeling, we washed out the exogenous dye and continued to acquire fluorescence images. To address the rate of fluorescence change during labeling and washout, we pooled the total cell fluorescence at each timepoint and quantified the mean total and volume-normalized fluorescence across cells (**Figure 6B**). During labeling, the volume-normalized fluorescence initially increased rapidly and then started to plateau; by 10 min, cells reached ∼50% of the labeling at 30 min, confirming rapid labeling of all three probes. During washout, total fluorescence was gradually lost (**Figure 6B**). We fit the washout dynamics to an exponential *I*(*t*) = *I*_0_ + *I*_1_*e*^−*γt*^. Washout of DMN-Tre labeling was slow, likely because it depends on mycomembrane turnover (**Figure 6Bi, Figure S2A**). Compared to DMN-Tre, cells labeld with 3HC-3-Tre or 3HC-2-Tre exhibited a rate of fluorescence loss *γ* that was 2.3- and 5.9-fold higher than DMN-Tre, respectively (**Figure 6Bii,iii, Figure S2B,C**), suggesting additional labeling mechanisms beyond trehalose synthesis. In particular, the washout time scale for 3HC-2-Tre was ln 2/ *γ* = 7.4 min, reflecting trehalose-independent transient binding and/or turn-on. Nonetheless, we confirmed that 3HC-3 and 3HC-3-Glc do not label Msmeg cells (**Figure S2E,F**), indicating the specificity of 3HC-3-Tre for trehalose. Taken together, these data demonstrate 3HC-3-Tre, but not 3HC-2-Tre, is an excellent candidate for the rapid detection of mycobacteria. Moving forward, we focused our efforts on the 3HC-3-Tre probe.

### 3HC-3-Tre labeling of Msmeg cells is much brighter than DMN-Tre and selective for Actinobacteria

Our next goal was to assess the specificity of 3HC-3-Tre labeling of mycobacteria compared with bacterial species that do not incorporate trehalose into their cell envelopes (**Figure 7**). We analyzed Msmeg cells incubated with 1, 10, or 100 µM of 3HC-3-Tre or 3HC-3-Glc (as a negative control) for 1 hour at 37 °C. We found that 100 µM 3HC-3-Tre-labeled cells were 100-fold brighter compared to background (**Figure 7A**). Importantly, in the same conditions, the fluorescence intensity of 3HC-3-Tre-labeled cells was 10-fold greater than that of DMN-Tre-labeled cells (**Figure 7B**), confirming that 3HC-3-Tre is a much brighter option than DMN-Tre. We previously reported that DMN-Tre selectively labeled organisms within the Actinobacteria suborder (16,19). To test whether a similar labeling pattern persisted for 3HC-3-Tre, we labeled Msmeg, *Corynebacterium glutamicum* (Cg), *Bacillus subtilis* (Bs), *Escherichia coli* (Ec) and *Staphylococcus aureus* (Sa) with 100 µM 3HC-3-Tre for 1 hour and analyzed the samples by microscopy and flow cytometry (**Figure 7C,D**). Similar to DMN-Tre, we observed bright labeling of Msmeg and Cg cells, consistent with both of these organisms utilizing trehalose in their cell envelopes. Although not detected by microscopy, there was minimal but significant fluorescence from labeled Bs, Ec, and Sa cells compared to the no-dye control using flow cytometry (**Figure 7C**), perhaps reflecting a trehalose-independent pathway that led to faster washout of 3HC-3-Tre than DMN-Tre (**Figure 6B**).

**Figure 7.**
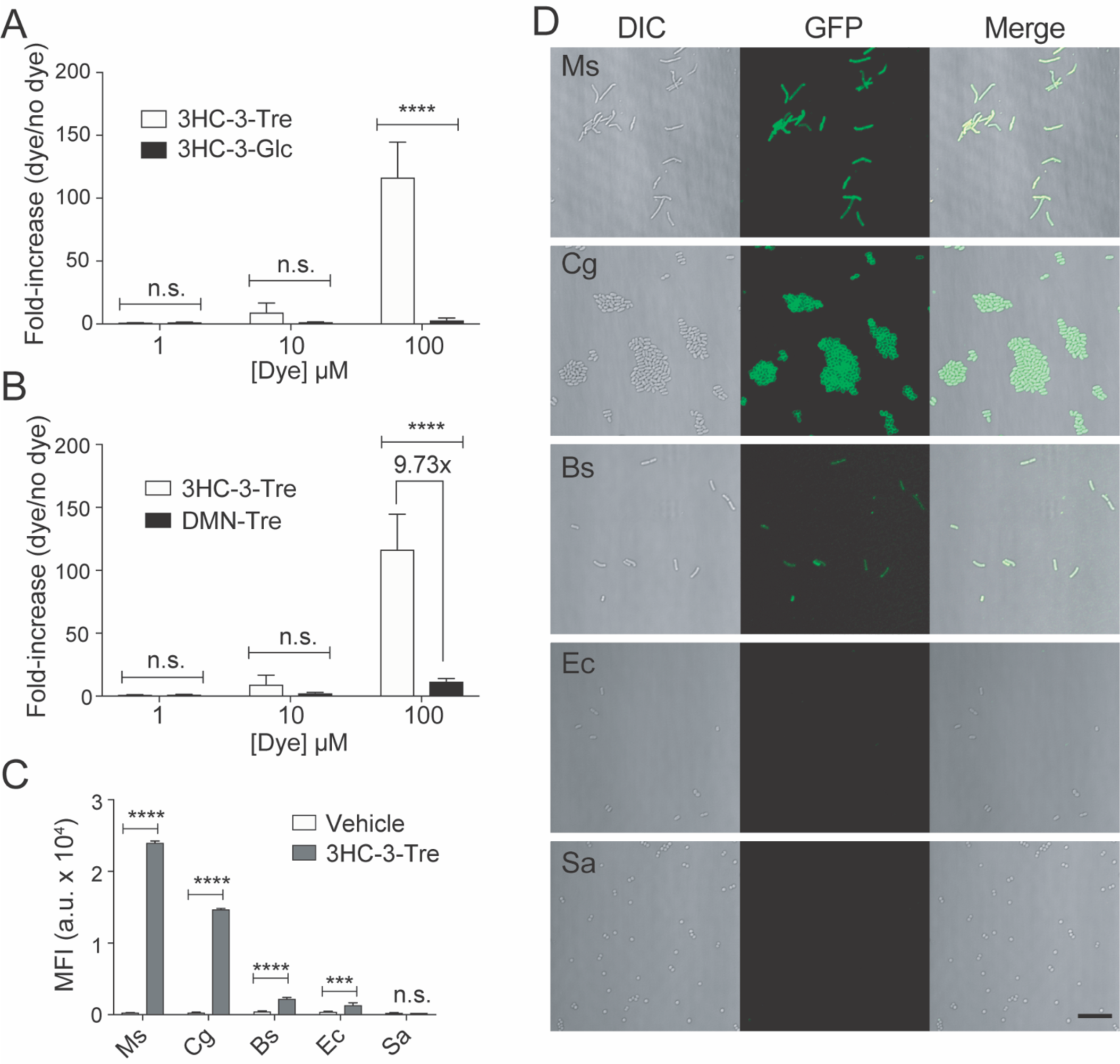
Specific labeling of live mycobacteria and corynebacteria with 3HC-3-Tre. **(A,B**) Flow cytometry analysis of Msmeg cells incubated for 1 h at 37 °C with (**A**) 100 µM 3HC-3-Tre or 3HC-3-Glc, (**B**) 100 µM 3HC-3-Tre or DMN-Tre. (**C,D**) Flow cytometry (**C**) and no-wash microscopy (**D**) analyses of Msmeg (Ms), *C. glutamicum* (Cg), *B. subtilis* (Bs), *E. coli* (Ec), and *S. aureus* (Sa) cells incubated for 1 h at 37 °C with 100 μM 3HC-3-Tre. 3HC-3-Tre labeling was specific to Msmeg and Cg. Data are means ± SEM from at least two independent experiments. Data were analyzed by two-way analysis of variance (ANOVA) test (*: *p*<0.05, **: *p*<0.01, ***: *p*<0.001, ****: *p*<0.0001, ns: not significant). Scale bar: 10 µm.

Finally, we assessed labeling of Mtb with 3HC-3-Tre (**Figure 8**). We labeled Mtb strain H37Ra cells with 100 µM 3HC-3-Tre for 1, 2, or 24 hours and then performed microscopy (**Figure 8A**). Similar to DMN-Tre, we observed labeling within 1 hour, with little discernible increase in fluorescence intensity over additional time (**Figure 8A**). By flow cytometry, we were able to detect significant fluorescence over background within 10 minutes of Mtb H37Ra labeling with 3HC-3-Tre (**Figure 8B**). In the same conditions, with DMN-Tre we required a minimum of 60 minutes to accurately detect significant fluorescence over background (19), likely due to the weak brightness of the fluorophore.

**Figure 8.**
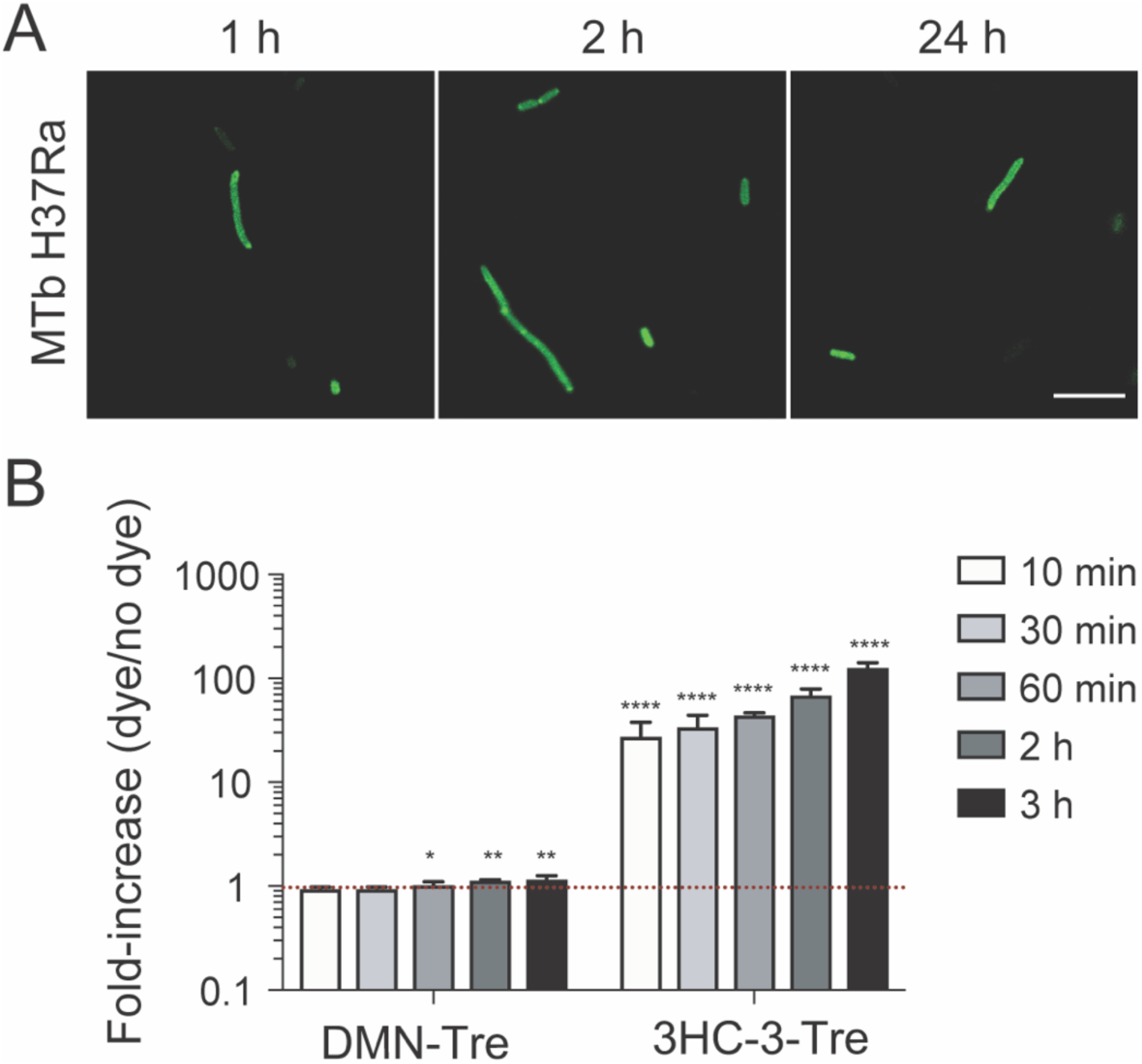
Mtb cells labeled with 3HC-3-Tre exhibit increased fluorescence intensities and can be detected within 10 minutes by flow cytometry. (**A**) Microscopy analysis of H37Ra Mtb cells labeled with 100 µM 3HC-3-Tre for 1, 2, or 24 h, followed by 4% paraformaldehyde fixation and fluorescence imaging. Scale bar: 10 µm. (**B**) Flow cytometry analysis of Mtb cells incubated for the indicated times with 100 µM DMN-Tre or 3HC-3-Tre, followed by 4% paraformaldehyde fixation. Ten minutes was sufficient for nearly complete labeling. Data are means ± SEM from at least two independent experiments. Data were analyzed by two-way analysis of variance (ANOVA) test (*: *p*<0.05, **: *p*<0.01, ***: *p*<0.001, ****: *p*<0.0001, ns: not significant).

## DISCUSSION

Tuberculosis (TB) remains a major global health threat (1). As it stands, poor detection methods have contributed to millions of missed TB diagnoses in high-burden, endemic countries (1,2). Due to the lack of accurate detection tests at the point-of-care, TB transmission rates have been sustained in low-income countries. In this study, we report on a solvatochromic dye trehalose conjugate that can swiftly detect mycobacteria. 3HC-3-Tre is a robust fluorogenic probe that specifically labels Actinobacteria such as Mtb within 10 minutes and can be imaged without any wash steps. In addition to its utility for research, this set of attributes may enable the rapid detection of Mtb in sputum samples in low-resource settings.

In future work, it may be possible to develop further red-shifted probes such as Nile-red dyes to minimize background fluorescence and maximize signal-to-background ratio, while maintaining minimal perturbations to mycobacteria. Furthermore, it is easily conceivable to synthesize trehalose moieties conjugated to an assortment of colors that span the fluorescence wavelength range to achieve multimodal labeling of mycobacteria. In addition to trehalose, the dyes could be coupled with other types of bacteria-specific molecules (sugars or amino acids) that would permit visualization of a subject of interest in various biological systems.

Beyond the performance of the detection methods, TB diagnoses in low-income environments depend on many factors, such as stability, accessibility, and affordability (2). The solvatochromic dyes are highly stable at room temperature even in temperate environments, can potentially be coupled with a portable florescence detection device, and a reasonable estimate of cost would be less than 40 cents per test. Because of these unique attributes, it is conceivable to use these reagents as mobile biomarkers for TB screening in resource-poor, remote environments. Thus, these novel probes can be used in service of both the scientific community to uncover mycobacterial metabolism, and also the clinical community at large for TB detection in high-burden environments.

## METHODS

### Fluorescence spectra procedures

One microliter (μL) of each dye-Tre conjugate (10 mM in water) was added to 1 mL of water or 99.9%, 99%, 95%, 90%, 75%, or 50% 1,4-dioxane in 1 cm x 0.4 cm quartz cuvettes (Starna Cells, Inc. 9F-G-10). Fluorescence data were acquired on a Photon Technology International Quanta Master 4 L-format scanning spectrofluorometer equipped with an LPS-220B 75-W xenon lamp and power supply, an A-1010B lamp housing with an integrated igniter, a switchable 814 photon-counting/analog photomultiplier detection unit, and an MD5020 motor driver. In the associated FelixGX software (v. 4.3.4.2010.6904), spectra were acquired using standard emission scan settings with the exception of the lamp slit widths, which were all set to 1 nm. Compounds were excited at 405, 488, 532, or 561 nm and emission intensity was monitored over 415-600 nm, 500-700 nm, 545-700 nm, or 575-750 nm, respectively. Data were exported as a text file and processed in Excel. Prism 7 (GraphPad) was used to create the figures from the final data.

### General procedures for bacterial culture inoculation

Cultures of *Mycobacterium smegmatis* (Msmeg), *Mycobacterium marinum* (Mmar), *Corynebacterium glutamicum* (Cg), *Bacillus subtilis* (Bs), *Escherichia coli* (Ec), and *Staphylococcus aureus* (Sa) were grown as described previously (6). Briefly, Msmeg single colonies were inoculated in BD Difco Middlebrook 7H9 media (supplemented with 10% (v/v) oleate-albumin-dextrose-catalase (OADC) enrichment, 0.5% (v/v) glycerol, and 0.5% (w/v) Tween 80) and incubated at 37 °C overnight. For Cg, Bs, Ec, and Sa, single colonies were inoculated in LB medium. Cg cultures were incubated at 30 °C, and Bs, Ec, and Sa cultures were incubated at 37 °C, all overnight. A 1-mL aliquot of Mmar (stored as a frozen stock in 50% glycerol/50% 7H9 medium) was thawed, then pelleted at 3,300*g* for 3 min. Cells were resuspended in 7H9 media supplemented with 10% (v/v) OADC enrichment, 0.5% (v/v) glycerol, and 0.5% (w/v) Tween 80, and incubated at 33 °C overnight.

### Metabolic labeling experiments

Experiments were performed as described previously (6). Briefly, overnight bacterial cultures were grown or diluted to an OD_600_ of 0.5 and aliquoted into Eppendorf tubes. The appropriate amount of stock dye (1 or 10 mM) was added to the aliquots to reach the indicated final concentration. Control samples were treated identically, without the addition of any probes. The bacteria were incubated for 1 h at 37 °C (Msmeg, Bs, Ec, Sa) or 30 °C (Cg). At the end of the experiment, samples were analyzed by microscopy and/or flow cytometry as described below.

### Confocal microscopy

For no-wash imaging, a drop of sample (∼5 µL) was taken directly from the labeled culture. For wash imaging, cells were pelleted at 3300*g* for 3 min, washed twice with 1x Phosphate Buffered Saline (PBS) supplemented with 5% Tween80 (v/v), and resuspended in 300 μL 1x PBS. Subsequently, a drop of sample (∼5 µL) was spotted onto a 1% agarose pad on a microscope slide, allowed to dry, covered with a cover slip and sealed with nail polish. Microscopy was performed on a Nikon A1R confocal microscope equipped with a Plan Fluor 60x oil immersion (NA: 1.30) objective. Samples were excited with a 405-nm violet laser, 488-nm blue laser, or 561-nm green laser and imaged in the Aqua Amine (425-475 nm), FITC/GFP (500-550 nm), or RFP (570-620 nm) channels, respectively. NIS-Elements AR software (Nikon Inc.) and Fiji (ImageJ) v. 72 were used to process images. All image acquisition and processing were executed under identical conditions for control and test samples.

### Flow cytometry

Cells were pelleted at 3300*g* for 3 min, washed twice with 1x Dulbecco’s phosphate-buffered saline (DPBS; MT-21-030-CV, Thermo Fisher Scientific) supplemented with 5% Tween80 (v/v), and resuspended in 300 μL 1x DPBS. Fluorescence measurements were taken in 5-mL culture tubes (14-959A, Thermo Fisher Scientific) suitable for flow cytometry. Experiments were performed on a BD LSR II.UV instrument in the shared Fluorescence Activated Cell Sorting (FACS) Facility at Stanford University. The instrument, excitation wavelengths, and filter sets used are noted in each figure or figure caption. Data were obtained for 10,000 cells per sample, processed using FlowJo, and imported into Prism 7 (Graphpad) for statistical analysis.

### Single-cell time-lapse microscopy

Single-cell time-lapse imaging was achieved using a microfluidic flow cell (CellASIC, B04A) and a custom temperature-controlled microscope system. Samples of mid-log cultures (200 µL) were placed undiluted into loading wells. Wells containing 7H9 medium and 7H9 medium with dye were primed for 10 min at the target temperature under 5 psi. During the experiment, flow was set at 2 psi. Cells were imaged at 1-min time intervals using a Ti-Eclipse stand (Nikon Instruments) with a Plan Apo 100X DM Ph3 (NA: 1.45) (Nikon) objective and images were acquired with an iXon EM+ (Andor) camera. All cells were imaged in phase-contrast and TIRF illumination. Cells stained with dyes excited at peak GFP wavelength (488 nm; hydroxychromone dyes) were imaged under TIRF illumination to reduce background signal and photobleaching, for which an OBIS laser (Coherent) light path was guided by an optical fiber to a TIRF illuminator (Nikon) and focused on the sample. Temperature was maintained at 37 °C using a stage-top incubator (Haison) coupled to a heater-controller (Air-Therm). Timing and control of the system was accomplished through µManager v. 1.41 (29).

### Image analysis

Image stacks were imported into ImageJ (FIJI) for initial data processing. Individual isolated cells were selected and cropped as image hyperstacks to include both phase-contrast and fluorescence channels. Phase-contrast images were used for automated segmentation analysis using Morphometrics (30) in MATLAB (Mathworks), and outlines were overlaid on the corresponding fluorescence images for quantifying signal information. Further analyses were carried out using custom MATLAB scripts. Cellular intensity was normalized to the peak cellular intensity during labeling for dye constructs with high signal-to-background. The initial period of fluorescence intensity decay was quantified by nonlinear regression fitting to an exponential function (*I*(*t*) = *I*_0_ + *I*_1_*e*^-*γt*^). For dye constructs with no signal, cellular intensity was normalized by the mean signal-to-background of the cell to reflect the relative signal.

### Metabolic labeling of Mtb

Mtb cultures were grown via inoculation of a 1-mL frozen stock into 50 mL of Middlebrook 7H9 liquid medium supplemented with 10% (v/v) OADC enrichment (BBL Middlebrook OADC, 212351), 0.5% (v/v) glycerol, and 0.05% (w/v) Tween 80 (P1754, Sigma-Aldrich) in a roller bottle. Cells were grown to an OD_600_ of 0.5 to begin the experiments. Cells were incubated with 100 µM DMN-Tre or 10 µM 3HC3-Tre for the indicated times. Labeled cells were harvested by centrifugation (3000*g* for 10 min) and then fixed in an equal volume of 4% paraformaldehyde (cells were incubated at room temperature for 1 h, with occasional rotation of the tube to ensure sterilization of all internal surfaces before fluorescence and flow cytometry analysis)

### *S*tatistical analysis

Data are means ± SEM from at least two independent experiments. Unless otherwise specified, all data were analyzed using GraphPad Prism software’s ANOVA (analysis of variance) test, as specified in the figure legends.

## ACKNOWLEDGEMENTS

We thank M. Rajendram, D. M. Fox, and S. Banik for technical assistance and helpful discussions, and A. Iavarone [QB3/Chemistry Mass Spectrometry Facility, University of California (UC), Berkeley] for assistance with mass spectrometry. We also thank Aidan Pezacki for research assistance. Flow cytometry was performed in the shared FACS facility at Stanford University on equipment obtained with NIH S10 Shared Instrument grant S10RR027431-01. M.K. was supported by the Stanford University’s Diversifying Academia, Recruiting Excellence Fellowship, and NIH Predoctoral Fellowship F31AI129359. S.G.L.K. was supported in part by a National Science Foundation Graduate Research Fellowship (Grant No. DGE-1106400). K.C.H. is a Chan Zuckerberg Biohub Investigator. This research was supported by the Allen Discovery Center at Stanford on Systems Modeling of Infection (to K.C.H.), and grants from the Bill and Melinda Gates Foundation (OPP115061) and NIH (AI051622) (to C.R.B.).

