## Supplemental Information for "Towards *Mycobacterium tuberculosis* detection at the point-of-care: a brighter solvatochromic probe permits the detection of mycobacteria within minutes"

### I. Materials and Methods

### A. Synthetic procedures

#### A. 1. General

Reactions were performed in flame- or oven-dried glassware under an inert nitrogen atmosphere unless otherwise noted. Anhydrous solvents were either purchased or obtained by passing solvent through an activated alumina column via a Pure Process Technology Glass Contour Solvent Purification System. All reagents and solvents were used as received unless otherwise noted. Water was passed through a Milli-Q filtration system prior to use. Where noted, samples were concentrated *in vacuo* at 40 °C using a BÜCHI Rotavapor R-114 equipped with a BÜCHI B-480 heating bath and a Welch Self-Cleaning Dry Vacuum System (Model 2025) or an IKA RV 10 basic rotary evaporator equipped with an IKA HB 10 basic heating bath and a Welch Self-Cleaning Dry Vacuum System (Model 2025). If necessary, compounds were then further dried under high vacuum using an Edwards RV8 Two Stage Rotary Vane Pump or by lyophilization in a LABCONCO FreeZone 4.5Plus.

Thin layer chromatography was performed using SiliCycle SiliaPlate glass-backed silica gel plates containing a fluorescent indicator (Fisher Scientific 50964470). Plates were visualized using a UVGL-25 Compact UV Lamp, 254/365 nm, 4 W (P/N 95-0021-12). For flash column chromatography, the stationary phase was SiliCycle SiliaFlash P60 or Fisher Silica Gel Sorbent (230-400 Mesh, Grade 60) silica gel. For purifications involving preparative Reversed-Phase High-Performance Liquid Chromatography (RP-HPLC), the following conditions were used: the instrument consisted of an Agilent Technologies ProStar 325 UV-Vis detector, two PrepStar Solvent Delivery Modules, and a 440-LC Fraction Collector; the column was either a Varian Microsorb 100Å C18, 8 µm, 21.4 x 250 mm Dynamax preparative column (R0080220C8) equipped with a Microsorb 100 Å C18, 8 µm guard column (R0080220G8) or a Phenomenex Luna® 10 µm Phenyl-Hexyl 100 Å, 250 x 21.2 mm preparative column (00G-4325-P0-AX). For the C18 column, solvent A was 0.1% TFA in Milli-Q water and solvent B was 0.1% TFA in acetonitrile (MeCN). For the phenyl-hexyl column, solvent A was pure Milli-Q water and solvent B was methanol. The UVVis detector was used to monitor wavelengths at 210 and 254 or 550 nm. All pure compounds and stock solutions were stored at -20 °C. For Isolera™ Prime purifications, the following general conditions were applied: the instrument was a Biotage Isolera™ Prime with one channel, a single collection bed, and a 200-400 nm detector (model ISOPSV). Specific solvents, monitored wavelengths, and column types are noted in individual synthesis sections.

High resolution mass spectrometry (HRMS) data were acquired by ESI-LC/MS on a Waters Acquity UPLC and Thermo Exactive Orbitrap mass spectrometer at the Stanford University Mass Spectrometry facility.

### A. 2. Sugars

**6-bromo-6-deoxy-α,α-trehalose heptaacetate (6-Br-$Ac_7$-Tre).** In a 200-mL flame-dried round-bottom flask equipped with a stir bar, anhydrous DMF (100 mL) was heated to 85 °C, then anhydrous trehalose (4 g, 11.69 mmol) and recrystallized NBS (2.29 g, 12.9 mmol, 1.1 equiv.) were dissolved therein. On addition of the NBS, the solution changed to pale yellow, then orange yellow, then orange. After 5 min, $PPh_3$ (6.13 g, 23.37 mmol, 2 equiv.) was added, and the solution immediately turned a clear/champagne color. The mixture was allowed to stir for 48 h at room temperature (r.t.), then the DMF was removed *in vacuo* to yield a clear off-white oil. Anhydrous pyridine (100 mL) was added, then the resulting solution was cooled to 0 °C and acetic anhydride (~15.5 mL, 164 mmol, 2 equiv. relative to the 7 hydroxyl groups on 6-Br-Tre) was added fast dropwise. The yellow solution was allowed to stir for 48 h at r.t., over which time it became a clear orange-red solution. After removing the solvent *in vacuo*, the resulting orange oil was re-dissolved in ethyl acetate, washed twice with concentrated sodium bicarbonate (orange aqueous layer) and twice with brine (clear aqueous layer). The clear orange organic layer was dried over anhydrous sodium sulfate, filtered, and concentrated *in vacuo*. Product was purified by silica gel chromatography on an Isolera Prime (Biotage) equipped with a Zip KP-Sil 120 g column equilibrated with 5 column volumes (CV) of the starting solvent mixture. The flow rate was 100 mL/min, and all fractions were collected. The gradient was as follows: 5% EtOAc in DCM for 1 CV, 5-30% EtOAc in DCM over 10 CV, 30% EtOAc in DCM for 2 CV. Pure fractions were determined by TLC (4:1 DCM/EtOAc), visualizing first with UV to check for triphenylphosphine oxide side product then by 5% $H_2SO_4$ in MeOH and charring. Of the three sugar spots observed, the top one was dibrominated side product, the middle was product, and the bottom was peracetylated trehalose side product. Pure fractions were concentrated *in vacuo* to a white crystalline foam and lyophilized to yield 0.9727 g (11.9%) of hard white crystals.

$^1$H NMR (500 MHz, Chloroform-*d*) δ 5.47 (q, J = 10.0 Hz, 2H), 5.32 (t, J = 4.0 Hz, 2H), 5.13 (dd, J = 10.3, 3.9 Hz, 1H), 5.07 – 4.91 (m, 3H), 4.21 (dd, J = 12.4, 6.0 Hz, 1H), 4.10 (q, J = 7.1 Hz, 2H), 4.06 – 3.97 (m, 2H), 3.41 – 3.27 (m, 2H), 2.14 – 2.05 (m, 11H), 2.05 – 1.98 (m, 9H), 1.24 (t, J = 7.1 Hz, 1H).

$^{13}C\{^{1}H\}$ NMR (126 MHz, Chloroform-*d*) δ 170.67, 170.02, 169.96, 169.66, 169.63, 169.57, 92.27, 91.82, 71.18, 70.14, 69.89, 69.73, 69.63, 69.30, 68.56, 68.26, 61.79, 60.45, 30.48, 21.02, 20.74, 20.67, 20.63, 14.25.

HRMS (ESI) *m*/*z*: $[M + Na]^+$ calculated for $C_{26}H_{35}BrO_{17}Na$ 721.0950; found 721.0932.

**Methyl 6-iodo-6-deoxy-α-D-glucopyranoside triacetate (Me 6-I-$Ac_3$-Glc).** In a 500-mL flame-dried, two-necked round-bottom flask equipped with a stir bar, methyl α-D-glucoside (2.5 g, 12.9 mmol), $PPh_3$ (5.065 g, 19.3 mmol, 1.5 equiv.), and imidazole (1.753 g, 25.7 mmol, 2 equiv.) were combined and refluxed in anhydrous THF (100 mL). A solution of iodine (I2; 4.902 g, 19.3 mmol, 1.5 equiv.) in anhydrous THF (25 mL) was added fast dropwise and the reaction was allowed to reflux a further 2 h. The reaction was allowed to cool to r.t., then the solution was vacuum-filtered to remove the white precipitate (imidazole salt), transferred to a 500-mL round-bottom flask, and concentrated *in vacuo* to a slightly cloudy viscous oil. The oil was dissolved in anhydrous pyridine (100 mL) and cooled to 0 °C, then acetic anhydride (7.3 mL, 77.2 mmol, 2 equiv. relative to the 3 free hydroxyl groups) was added fast dropwise. The reaction was allowed to stir overnight at r.t. under nitrogen gas then concentrated *in vacuo* to a white solid/yellow oil. Product was purified by silica gel chromatography on an Isolera Prime (Biotage) equipped with a Zip KP-Sil 120 g column equilibrated with 3 CV of the starting solvent mixture. The flow rate was 100 mL/min, the 254/280 nm wavelengths were monitored (iodine absorbs at 254 nm), and peaks that absorbed above 25 mAu were collected. The gradient was as follows: 10% EtOAc in hexanes for 2 CV, 10-65% EtOAc in hexanes over 6 CV, 65-100% EtOAc in hexanes over 1 CV, and 100% EtOAc for 2.5 CV. Pure fractions were determined by TLC (1:1 hexanes/EtOAc), visualizing first with UV then by 5% $H_2SO_4$ in MeOH and charring. Pure fractions were combined and concentrated *in vacuo* then recrystallized from acetone and hexanes. After high vacuum, 3.3066 g (59.7%) of white fluffy crystals were obtained.

$^{1}H$ NMR (500 MHz, Chloroform-*d*) δ 5.45 (dd, J = 10.2, 9.3 Hz, 1H), 4.94 (d, J = 3.7 Hz, 1H), 4.88 – 4.83 (m, 2H), 3.77 (ddd, J = 10.2, 8.3, 2.5 Hz, 1H), 3.46 (s, 3H), 3.28 (dd, J = 10.9, 2.5 Hz, 1H), 3.12 (dd, J = 10.9, 8.3 Hz, 1H), 2.06 (s, 3H), 2.04 (s, 3H), 1.99 (s, 3H).

$^{13}C\{^{1}H\}$ NMR (126 MHz, Chloroform-*d*) δ 170.34, 170.26, 169.89, 96.91, 72.68, 71.12, 69.87, 68.84, 55.99, 20.98, 20.92, 3.93.

HRMS (ESI) *m*/*z*: $[M + Na]^+$ calculated for $C_{13}H_{19}IO_8Na$ 453.0017; found 453.0010.

### A. 3. Dyes

The aldehyde precursors of 3HC-3 and 3HC-2 were synthesized following previously reported protocols (1,2). Each was then converted to the 3-hydroxychromone using procedures modified from (3) as described below.

***3HC-3***. In a flame-dried three-necked 25-mL roundbottom flask equipped with a stir bar, 7 diethylamino-9,9-dimethylfluorene-2-carbaldehyde (50 mg, 0.17 mmol) was dissolved in anhydrous MeOH (5 mL). 2'-hydroxyacetophenone (23 μL, 0.193 mmol, 1.14 equiv.) and crushed NaOH (20.4 mg, 0.51 mmol, 3 equiv.) were added and the bright yellow reaction was refluxed overnight at 75 °C. Due to the small volume, the reaction dried to a red-orange oil overnight; 5 mL of anhydrous MeOH was added to re-dissolve it and the reaction was allowed to reflux a further 2 h. After checking that no more product had formed by TLC (9:1 hexanes/EtOAc, visualized by UV), the orange solution was removed from heat and allowed to cool completely to r.t. over approximately 1 h. Then, 0.5 M NaOH (1.02 mL) was added slow dropwise while stirring thoroughly, making sure that no solid crashed out of the reaction mixture. To the resulting clear red solution was added 30% hydrogen peroxide (70 μL, 0.7 mmol, 4.1 equiv.) dropwise. The reaction was covered with foil and allowed to stir under nitrogen at r.t. overnight. The reaction mixture was directly purified by reversed-phase C18 chromatography on an Isolera Prime (Biotage) equipped with a SNAP Ultra 60 g column equilibrated with 5 CV of the starting solvent mixture. The flow rate was 75 mL/min, the 210/254 nm wavelengths were monitored, and peaks that absorbed above 5 mAu were collected. The column was visually monitored for elution of colored bands – the solvent system was held at 72% MeCN/water (no TFA) until a pale orange band (yellow fractions) and a red-orange band (orange and red fractions) had mostly eluted (~26 CV), then the %MeCN was increased to 100% over 2 CV and held at 100% for 5 CV. Product eluted around CV 27~30 (a smaller peak very soon after/overlapping with the tail end of a large peak). Pure fractions (determined by LC-MS) were combined and concentrated *in vacuo*, then the product was dried under high vacuum overnight to yield 6.8 mg (9.7%) of an orange solid.

$^{1}$H NMR (500 MHz, Acetone-*d*) δ 8.39 (d, J = 1.7 Hz, 1H), 8.29 (dd, J = 8.2, 1.7 Hz, 1H), 8.18 (dd, J = 8.0, 1.6 Hz, 1H), 7.85 – 7.74 (m, 3H), 7.67 (d, J = 8.5 Hz, 1H), 7.51 – 7.46 (m, 1H), 6.92 (d, J = 2.4 Hz, 1H), 6.74 (dd, J = 8.5, 2.4 Hz, 1H), 3.51 (q, J = 7.1 Hz, 4H), 1.53 (s, 6H), 1.21 (t, J = 7.0 Hz, 6H).

HRMS (ESI) *m*/*z*: $[M + H]^+$ calculated for $C_{28}H_{28}NO_3$ 426.2064; found 426.2070.

***3HC-2***. A flame-dried 15-mL two-necked round-bottom flask equipped with a stir bar was charged with a solution of 6-diethylaminobenzofuran-2-carbaldehyde (41.8 mg, 0.19 mmol) in anhydrous MeOH (1.16 mL). 2'-hydroxyacetophenone (23 µL, 0.19 mmol) and crushed NaOH (23.1 mg, 0.58 mmol, 3 equiv.) were added and the reaction was refluxed overnight at 75 °C. Due to the small volume, the reaction dried to a red-orange solid overnight; 5 mL of MeOH was added to re-dissolve it and the reaction was allowed to reflux a further 2 h. After checking that no more product had formed by TLC (2:1 hexanes/EtOAc, visualized by UV), the orange solution was removed from heat and allowed to cool completely to r.t. over approximately 1 h. Then, 0.5 M NaOH (1.16 mL) was added slow dropwise while stirring thoroughly, making sure that no solid crashed out of the reaction mixture. To the resulting clear red solution was added 35% hydrogen peroxide (79 µL, 0.8 mmol, 4.1 equiv.) dropwise. The reaction was covered with foil and allowed to stir for 3 h at r.t. under nitrogen. TLC and LC-MS confirmed completion of the reaction, which was then poured into water. The MeOH was removed *in vacuo* and the remaining aqueous solution was extracted several times with EtOAc. The combined EtOAc layers (clear orange solution tinted green) were dried over anhydrous sodium sulfate, filtered, concentrated *in vacuo*, and dried under high vacuum overnight to yield 49.2 mg (73.2%) of an orange-red solid. [If necessary, the product was purified by silica gel chromatography on an Isolera Prime (Biotage) equipped with a SNAP KP-Sil 50 g column equilibrated with 5 CV of the starting solvent mixture. The flow rate was 100 mL/min, the 254/280 nm wavelengths were monitored, and peaks that absorbed above 40 mAu were collected. The gradient was as follows: 8% EtOAc/hexanes for 1 CV, 8-66% EtOAc/hexanes over 10 CV, 66% EtOAc/hexanes for 2 CV. Pure fractions were determined by TLC (4:1 hexanes/EtOAc, visualized with UV) and LC-MS, concentrated *in vacuo*, and dried under high vacuum overnight.]

HRMS (ESI) *m*/*z*: $[M + H]^+$ calculated for $C_{21}H_{19}NO_4$ 350.1387; found 350.1387.

$^1$H NMR (500 MHz, DMF-$d_7$) δ 10.36 (s, 1H), 8.97 (s, 1H), 8.17 (dd, *J* = 8.0, 1.6 Hz, 1H), 7.85 (ddd, *J* = 8.6, 6.9, 1.7 Hz, 1H), 7.78 (dd, *J* = 8.6, 1.1 Hz, 1H), 7.71 (d, *J* = 0.9 Hz, 1H), 7.62 (d, *J* = 8.6 Hz, 1H), 7.52 (ddd, *J* = 8.0, 6.9, 1.1 Hz, 1H), 6.94 – 6.86 (m, 2H), 6.83 (dd, *J* = 5.8, 3.6 Hz, 1H), 6.65 (dd, *J* = 5.9, 3.5 Hz, 1H), 6.52 – 6.45 (m, 1H), 3.54 (d, *J* = 7.0 Hz, 4H), 1.22 (t, *J* = 7.0 Hz, 6H).

$^{13}$C NMR (126 MHz, DMF-$d_7$) δ 177.06, 171.64, 158.03, 154.91, 146.10, 143.58, 138.45, 133.78, 125.16, 124.90, 122.89, 122.63, 119.67, 118.51, 117.80, 117.34, 115.87, 112.61, 110.84, 107.75, 95.80, 92.81, 44.82, 12.39.

### A. 4. Conjugation of dyes to trehalose or glucose

This procedure was adapted from (4). Briefly, in a flame-dried 10-mL pear-shaped flask, dye (0.02 mmol), anhydrous potassium carbonate (0.0354 mmol, 1.77 equiv.), and anhydrous DMF (~1 mL or enough to fully dissolve the dye, whichever is greater) were combined and stirred for 1 h at r.t. Then, 6-Br-$Ac_7$-Tre or Me 6-I-Ac3-Glc (0.06 mmol, 3 equiv.) was added, and the reaction was allowed to stir 48 h at 80 °C, protected from light. The reaction was allowed to completely cool to r.t., then anhydrous 0.5 M NaOMe in MeOH (~1 mL) was added directly to the mixture. After stirring for 1.5 h at r.t., the reaction was directly purified by reversed-phase C18 chromatography on an Isolera Prime (Biotage) equipped with a SNAP Ultra C18 60 g column (75 mL/min) or a SNAP Ultra C18 30 g column (50 mL/min) equilibrated with 5-7 CV of the starting solvent mixture. No TFA was added. The gradient was modified on the fly: when a 210/254 nm peak appeared or colored bands on the column began to move, the gradient was held at that solvent ratio until the compound eluted. The method was set to collect peaks absorbing above 3 mAu (210/254 nm), but in practice the user told the machine when to collect based on whether a colored band was eluting. The basic gradient was 100% water for 3 CV, 0-100% MeCN/water over 15 CV, 100% MeCN. Pure fractions were determined by LC-MS, combined, and concentrated *in vacuo*, then resuspended in Milli-Q water and lyophilized overnight.

**3HC-3-Tre**. 3HC-3 (5 mg, 0.012 mmol) yielded 3.291 mg (37.4% over three steps) of an orange powder.

$^1$H NMR (500 MHz, DMSO-d6) δ 8.32 – 8.22 (m, 2H), 7.88 – 7.80 (m, 2H), 7.68 (d, J = 8.5 Hz, 1H), 6.84 (d, J = 2.3 Hz, 1H), 6.68 (dd, J = 8.6, 2.4 Hz, 1H), 5.10 (d, J = 5.0 Hz, 1H), 4.95 (d, J = 3.6 Hz, 1H), 4.88 (dd, J = 10.9, 4.2 Hz, 2H), 4.79 (dd, J = 5.1, 2.3 Hz, 2H), 4.70 (dd, J = 6.1, 1.9 Hz, 2H), 4.39 – 4.31 (m, 2H), 4.18 (dd, J = 10.6, 5.1 Hz, 1H), 4.08 – 4.01 (m, 1H), 3.69 (ddd, J = 9.9, 4.8, 2.4 Hz, 1H), 3.65 – 3.48 (m, 3H), 3.45 (p, J = 7.6, 7.0 Hz, 4H), 3.35 (s, 31H), 3.27 – 3.19 (m, 2H), 3.19 – 3.10 (m, 1H), 1.48 (d, J = 7.5 Hz, 6H), 1.15 (t, J = 7.0 Hz, 6H).

$^{13}$C NMR (126 MHz, DMSO-d6) δ 174.67, 157.21, 156.24, 155.30, 153.11, 149.01, 143.36, 140.14, 134.60, 128.79, 126.97, 125.66, 125.55, 124.05, 123.00, 122.81, 119.15, 118.81, 111.25, 105.96, 94.27, 73.52, 73.21, 73.16, 72.29, 72.24, 71.77, 70.86, 70.72, 67.05, 61.46, 47.15, 44.70, 27.74, 27.70, 13.20.

HRMS (ESI) *m/z*: $[M + H]^+$ calculated for $C_{40}H_{48}NO_{13}$ 750.3120; found 750.3115.

**3HC-3-Glc.** 3HC-3 (3 mg, 0.007 mmol) yielded 0.338 mg (8% over three steps) of a yellow-orange powder.

$^1$H NMR (500 MHz, acetone-d6) δ 8.56 (s, 1H), 8.21 (t, J = 9.3 Hz, 1H), 7.90 – 7.75 (m, 3H), 7.70 (d, J = 8.5 Hz, 1H), 7.55 – 7.50 (m, 1H), 7.21 (s, 1H), 6.93 (s, 1H), 6.75 (d, J = 8.5 Hz, 1H), 5.26 (s, 1H), 5.11 (s, 1H), 4.76 (s, 1H), 4.35 (s, 1H), 4.26 (d, J = 10.7 Hz, 1H), 4.00 (s, 1H), 3.82 (d, J = 9.0 Hz, 1H), 3.77 (s, 1H), 3.71 (s, 1H), 3.59 (d, J = 7.4 Hz, 1H), 3.52 (q, J = 7.7, 7.3 Hz, 4H), 3.33 (d, J = 24.9 Hz, 3H), 1.56 (d, J = 19.1 Hz, 6H), 1.22 (t, J = 7.2 Hz, 6H).

HRMS (ESI) *m/z*: $[M + H]^+$ calculated for $C_{35}H_{40}NO_8$ 602.2748; found 602.2760.

**3HC-2-Tre.** 3HC-2 (10 mg, 0.031 mmol) yielded 1.672 mg (8.7% over three steps) of an orange fluffy solid.

$^1$H NMR (500 MHz, Methanol-d4) δ 8.19 (d, J = 8.0 Hz, 1H), 7.99 (s, 1H), 7.84 – 7.78 (m, 1H), 7.72 (d, J = 8.3 Hz, 1H), 7.55 (d, J = 9.2 Hz, 1H), 7.49 (s, 1H), 6.85 (s, 2H), 5.27 (s, 1H), 5.17 (s, 1H), 4.46 (d, J = 14.9 Hz, 2H), 4.19 (d, J = 9.0 Hz, 1H), 3.90 (d, J = 10.4 Hz, 2H), 3.81 (d, J = 12.1 Hz, 2H), 3.73 (t, J = 8.5 Hz, 2H), 3.61 (d, J = 9.5 Hz, 1H), 3.54 – 3.44 (m, 6H), 1.27 – 1.21 (m, 6H).

HRMS (ESI) *m/z*: $[M + H]^+$ calculated for $C_{33}H_{40}NO_{14}$ 674.2443; found 674.2440.

**3HC-2-Glc.** 3-HC-2 (10 mg, 0.031 mmol) yielded 0.845 mg (5.2% over three steps) of an orange powder.

1H NMR (500 MHz, Methanol-d4) δ 8.19 (d, J = 8.0 Hz, 1H), 8.01 (d, J = 2.0 Hz, 1H), 7.81 (dd, J = 9.5, 7.5 Hz, 1H), 7.71 (d, J = 8.6 Hz, 1H), 7.53 – 7.47 (m, 3H), 7.29 (d, J = 3.6 Hz, 2H), 6.85 (d, J = 4.8 Hz, 2H), 4.80 (d, J = 3.8 Hz, 1H), 4.52 (d, J = 11.1 Hz, 1H), 4.47 (d, J = 5.0 Hz, 1H), 3.89 (s, 1H), 3.72 (t, J = 9.3 Hz, 1H), 3.61 (t, J = 9.5 Hz, 1H), 3.54 – 3.46 (m, 5H), 3.42 (s, 3H), 1.24 (t, J = 7.0 Hz, 6H).

HRMS (ESI) *m/z*: $[M + H]^+$ calculated for $C_{28}H_{32}NO_9$ 526.2072; found 526.2070.

DMN-Tre and DMN-Glc were on hand from a previous study (5).

### II. Supporting Schemes, Figures and Tables

#### A. Synthetic schemes

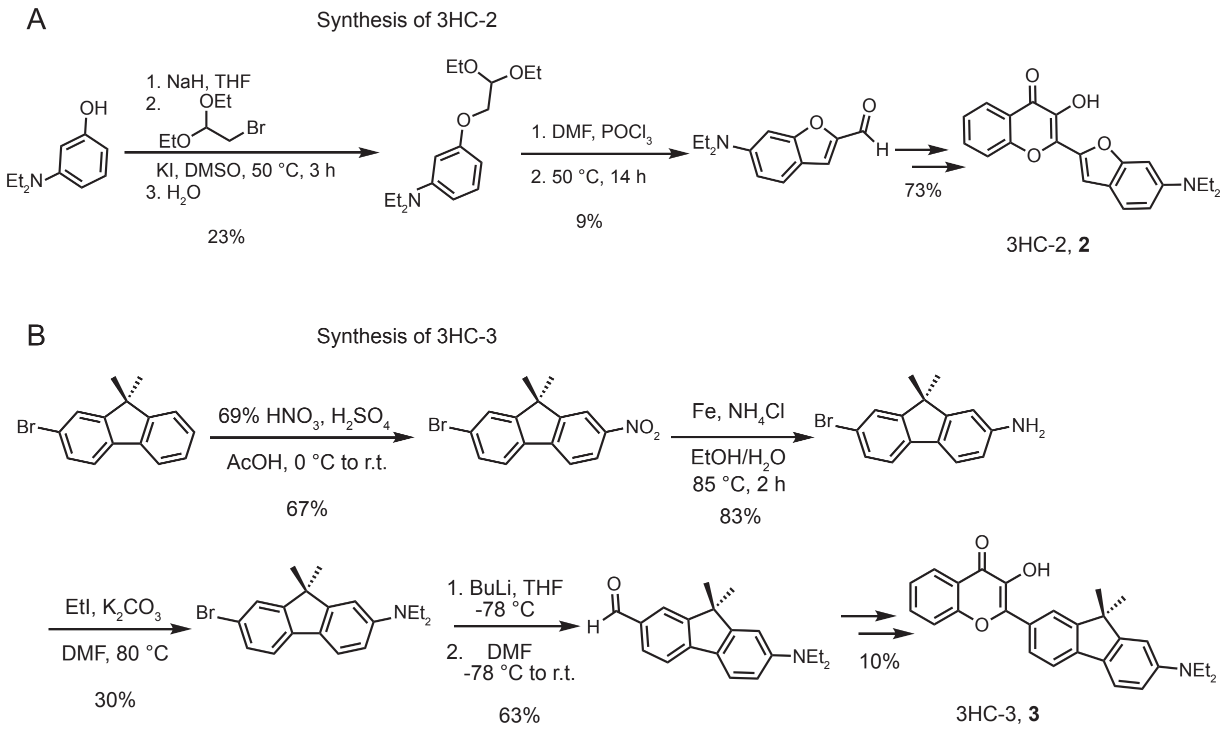

**Scheme S1. Synthesis of 3-hydroxychromone (3HC) dyes.**

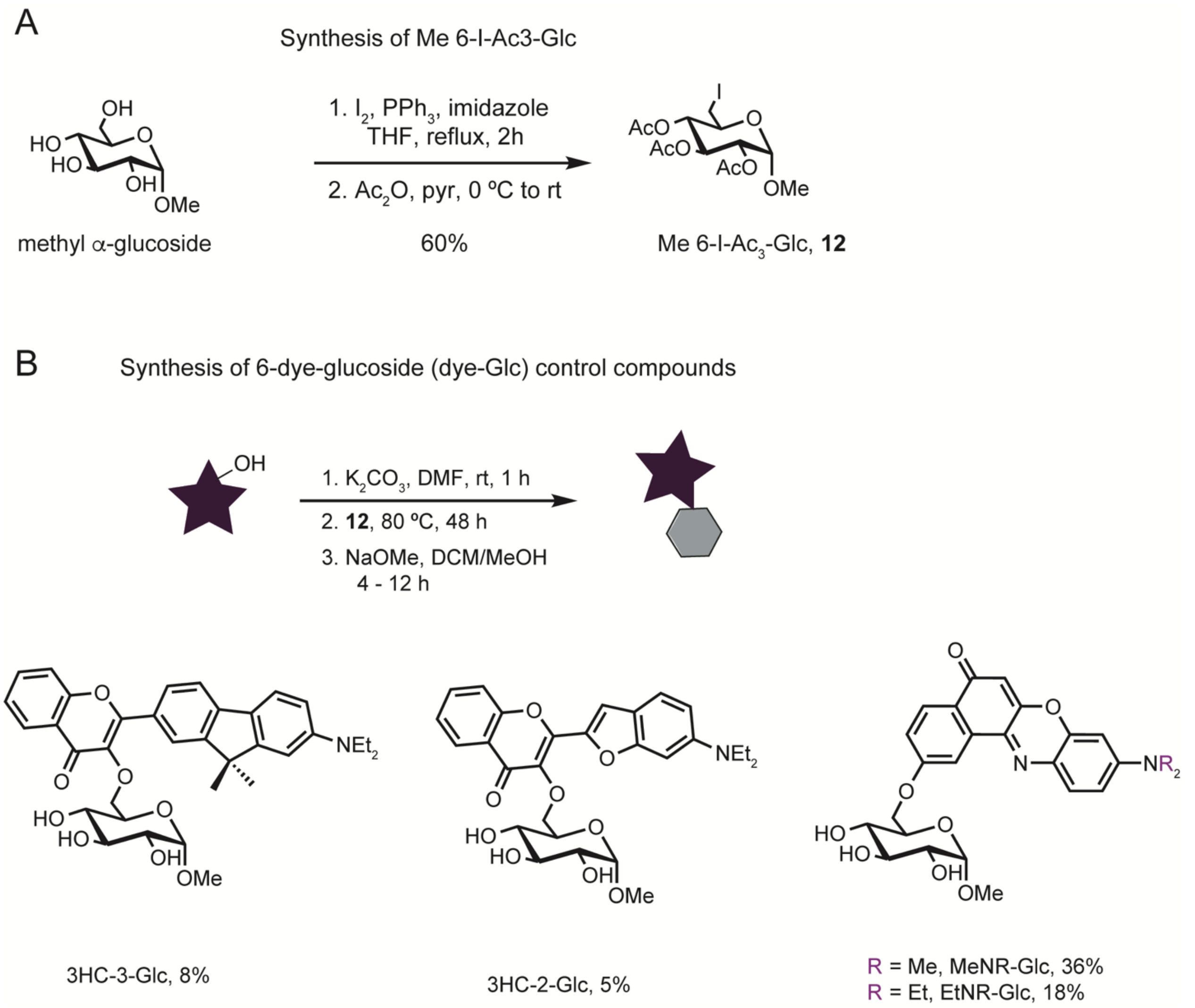

**Scheme S2: Synthesis of dye-glucose probes.**

### B. Supporting figures

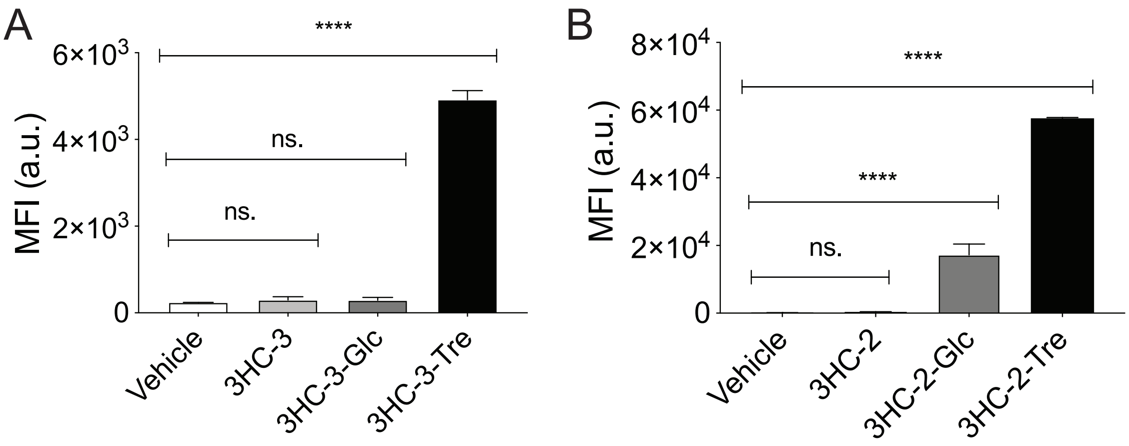

**Figure S1. Unlike 3HC-2-Tre, 3HC-3-Tre labeling is dependent on the trehalose moeity.** Flow cytometry analysis of Msmeg labeled with (**A**) 3HC-3, 3HC-3-Glc, or 3HC-3-Tre, or (**B**) 3HC-2, 3HC-2-Glc, or 3HC-2-Tre. Cells at $OD_{600}$=0.5 were incubated with the indicated dye trehalose probe concentrations for 1 h at 37 °C. MFI: Mean fluorescence intensity. Data are means ± SEM from at least two independent experiments. Data were analyzed by two-way analysis of variance (ANOVA) test (*: $p$<0. 05, **: $p$<0.01, ***: $p$< 0.001, ****: $p$<0.0001, ns: not significant).

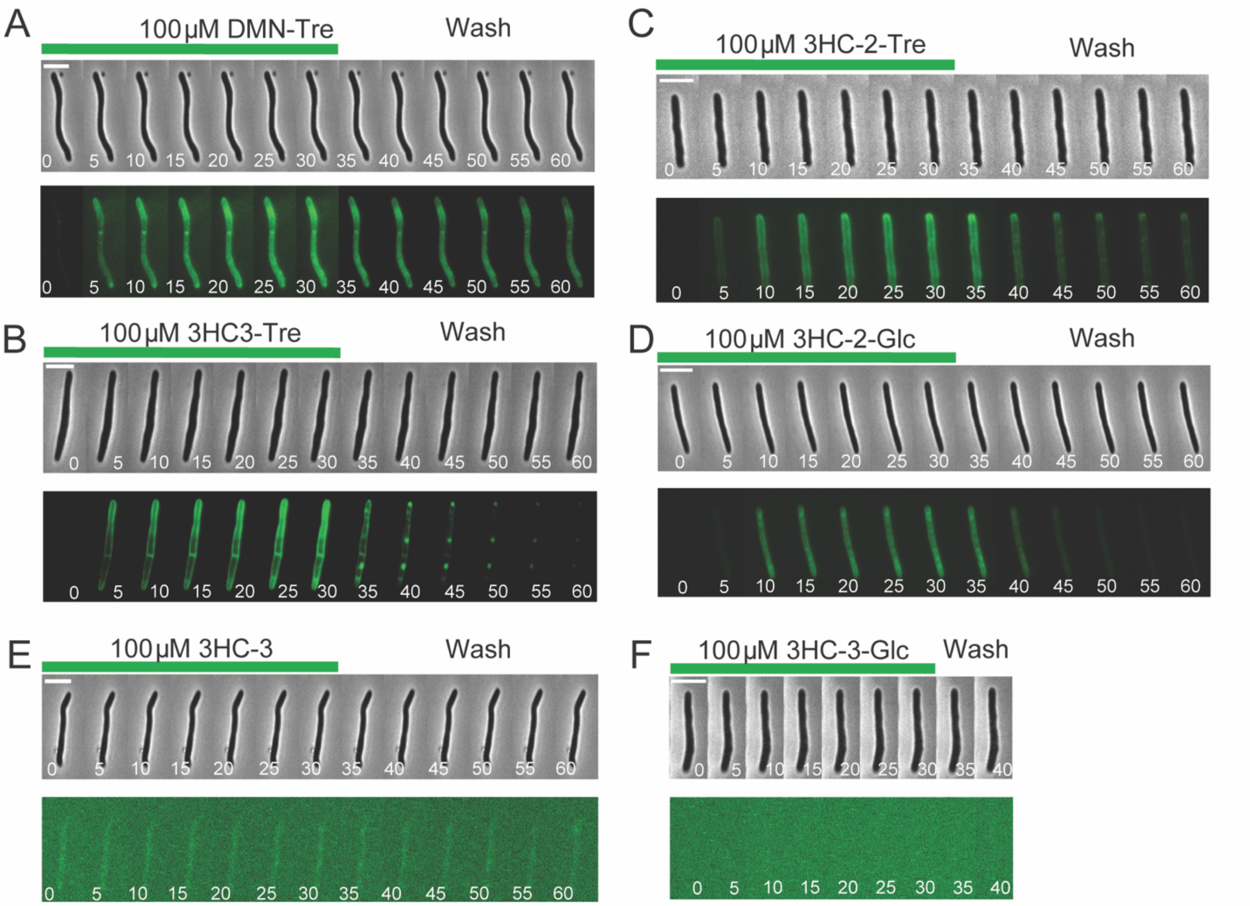

**Figure S2**. **Time-lapse epifluorescence microscopy of Msmeg labeling with solvatochromic trehalose probes**. Cells were treated with 100 μM (**A**) DMN-Tre, (**B**) 3HC-3-Tre, (**C**) 3HC-2-Tre, (**D**) 3HC-2-Glc, (**E**) 3HC-3 or (**F**) 3HC-3-Glc for 30 min, followed by 30 min of washout with liquid medium without dye.
